# Nucleoporin107 mediates female sexual differentiation via Dsx

**DOI:** 10.1101/2021.08.16.456487

**Authors:** Tikva Shore, Tgst Levi, Rachel Kalifa, Amatzia Dreifuss, Dina Rekler, Ariella Weinberg-Shukron, Yuval Nevo, Tzofia Bialistoky, Victoria Moyal, Merav Yaffa Gold, Shira Leebhoff, David Zangen, Girish Deshpande, Offer Gerlitz

## Abstract

We recently identified a missense mutation in Nucleoporin107 (Nup107; D447N) underlying XX-ovarian-dysgenesis, a rare disorder characterized by underdeveloped and dysfunctional ovaries. Modelling of the human mutation in Drosophila or specific knockdown of Nup107 in the gonadal soma resulted in ovarian-dysgenesis-like phenotypes. Transcriptomic analysis identified the somatic sex-determination gene doublesex (dsx) as a target of Nup107. Establishing Dsx as a primary relevant target of Nup107, either loss or gain of Dsx in the gonadal soma is sufficient to mimic or rescue the phenotypes induced by Nup107 loss. Importantly, the aberrant phenotypes induced by compromising either Nup107 or dsx are reminiscent of BMP signaling hyperactivation. Remarkably, in this context, the metalloprotease AdamTS-A, a transcriptional target of both Dsx and Nup107, is necessary for the calibration of BMP signaling. As modulation of BMP signaling is a conserved critical determinant of soma-germline interaction, the sex and tissue specific deployment of Dsx-F by Nup107 seems crucial for the maintenance of the homeostatic balance between the germ cells and somatic gonadal cells.

## Introduction

Germline-soma communication lies at the heart of proper gonad development, and thus is essential for formation and coalescence of the primitive embryonic gonad until generation of the adult gonad (*1–3*). Gonadogenesis also relies upon coordination between autonomous and non-autonomous cell mechanisms that direct correct specification, patterning, and subsequent morphogenesis (*4, 5*). Furthermore, the developmental program culminating in proper gonad formation must integrate both non-sex-specific ‘housekeeping’ functions and sex-specific signals to establish and maintain sexually dimorphic traits. Therefore, functional aberrations that affect individual molecular components, either sex-specific or non-sex-specific, result in various clinical disorders and infertility.

XX ovarian dysgenesis (XX-OD) is a rare, genetically heterogeneous disorder that is characterized by underdeveloped, dysfunctional ovaries, with subsequent lack of spontaneous pubertal development, primary amenorrhea, uterine hypoplasia, and hypergonadotropic hypogonadism (*6*). We recently identified a recessive missense mutation in the nucleoporin-107 (*Nup107*) gene (c.1339G>A, p.D447N) as the causative mutation for isolated XX-OD (without other developmental deficits) in five female cousins from a consanguineous family (*7*). All men in the family had normal pubertal development and those married have multiple children.

Nup107 is an essential component of the nuclear pore complex, enabling both active and passive transport in every nucleated cell. In light of this ubiquitous and vital function, the sex-specific and tissue-restricted nature of the *Nup107* XX-OD phenotype is highly intriguing. Taking into account the high conservation of the aspartic acid residue altered by the mutation, we used a *Drosophila* model to assess Nup107 function in human female gonadal development. Our previous studies demonstrated that RNAi-mediated knockdown (KD) of *Nup107* in somatic gonadal cells led to female-specific sterility due to defective oogenesis, while male flies developed normally and remained fertile (*7*). Furthermore, generating a *Drosophila* model of the human mutation by introducing an *RFP-Nup107^WT^* transgene in *Nup107*-null flies rescued both lethality and fertility of female flies; however, introduction of an *RFP-Nup107^D364N^* transgene recapitulating the familial XX-OD mutation, while rescuing lethality, resulted in severely reduced female fertility, ovariole disintegration, and extensive apoptosis (*7*).

A second mutation in *Nup107* (c.1063C>T, p.R355C) has since been independently identified as a cause of XX-OD (*8*). Notably, our analysis had predicted salt bridge interactions between the altered Aspartic Acid 447 in our study and the Arginine 355 altered in the second family, strengthening the notion that Nup107 performs an essential and conserved ovary-specific function during female gonadogenesis. Here, to uncover the molecular underpinnings of Nup107’s function, we analyzed the ‘loss of function’ phenotypes of *Nup107* at both the cellular and transcriptional levels.

## Results

### Nup107 mutant ovaries display an ovarian dysgenesis like phenotype

A closer inspection of *Nup107^D364N^* adult ovaries revealed that the female gonads displayed phenotypic traits closely resembling human ovarian dysgenesis. Specifically, the phenotypes ranged from rudimentary, small ovaries with fewer ovarioles to bilateral dysgenesis where both ovaries were completely absent (Figure. 1B-D). Approximately 33% of *Nup107^D364N^* ovaries were either under- or non-developed, in comparison to milder defects in only 6% and 5% in *Nup107^WT^* and *yw* control flies, respectively (Figure. 1A, G). We found similar ovarian defects when inserting the Nup107 mutation (1090G >A, p.D447N) into the Drosophila genome using CRISPR (*9*).

**Figure. 1.**
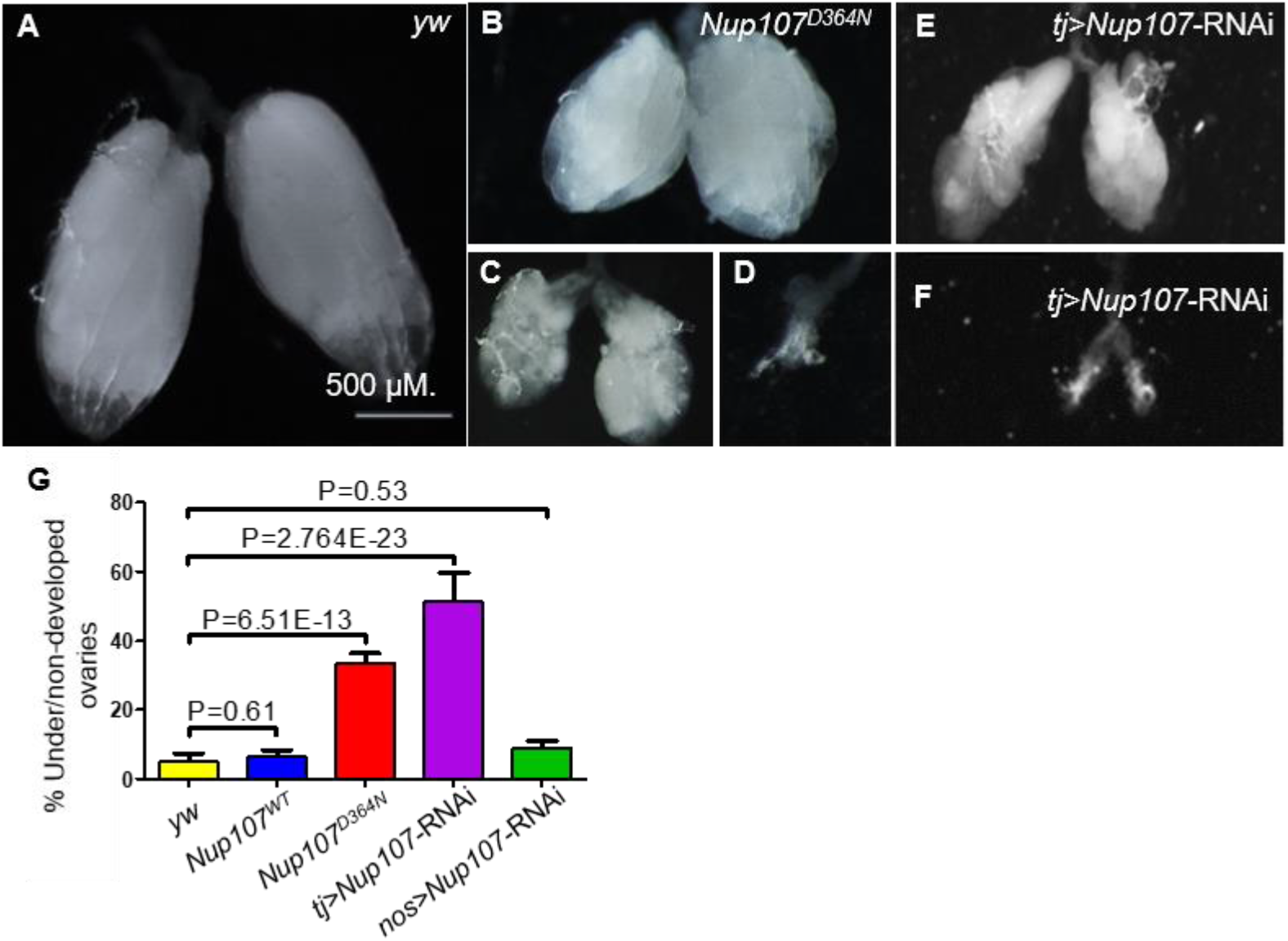
Phenotypic characterization of the ovaries compromised for *Nup107*. (**A**) *yw* control *Drosophila* ovaries. In contrast, *Nup107^D364N^* ovarian samples display a variety of aberrations, including (**B**) small or (**C**) shriveled ovaries, and (**D**) bilateral dysgenesis. Knockdown of Nup107 using *tj-Gal4* driver recapitulated the same phenotypes as *Nup107^D364N^*, with flies exhibiting (**E**) underdeveloped or (**F**) non-developed ovaries. (**G**) The percentages of under/non-developed ovaries in *yw* (n=196), *Nup107^WT^* (n=504), *Nup107^D364N^* (n=890), *tj>Nup107* (n=316), and *nos>Nup107* (n=64) flies.

To discern if Nup107 is necessary in the somatic or germline component of the gonad, or in both, we selectively inactivated *Nup107* function using *UAS-Nup107-RNAi* in combination with either a somatic gonadal specific driver *traffic jam–Gal4* (*tj-Gal4*) or a germ cell specific driver (*nanos* i.e., *nos-Gal4*). Interestingly, only soma specific knockdown of *Nup107* recapitulated the mutant phenotype, where 51% of adult ovaries were ‘dysgenic’ or under-developed (Figure. 1E-G). Supporting the conclusion that in this functional context Nup107 is required primarily in the soma, germline specific inactivation of Nup107 did not lead to significant phenotypic abnormalities compared to the control (9%, Figure 1G).

### Nup107 mutant ovaries display increased BMP signaling activity away from the germline stem cells (GSC) niche

To understand the functional underpinnings of the ovarian phenotypes induced by the D364N mutation we sought to analyze the Nup107 mutant ovaries carefully. The *Drosophila* ovary is made up of 16-20 ovarioles that function individually as egg production lines. The germarium, situated at the anterior tip of the ovariole, contains the somatic stem cell niche and adjacent Germline Stem Cells (GSCs) (*10, 11*). Typically, a GSC divides asymmetrically to generate another stem cell and a cystoblast. Cystoblasts divide and differentiate to form sixteen interconnected cystocytes, with the dynamically expanding fusome acting as the connecting link (Figure. 2A) (*12, 13*). Thus, GSCs can be readily identified by the presence of a round fusome, whereas their differentiated daughter cells display extended, branched fusomes. Ovary staining revealed that unlike *yw* and *Nup107^WT^* germaria, which contained 2-3 spherical fusomes, *Nup107^D364N^* germaria contained an average of 6 spherical fusomes per germarium (7% of *Nup107^WT^* vs. 34% of *Nup107^D364N^*; Figure. 2B-D). Similar results were seen in germaria where *Nup107* expression was knocked down (KD) using the somatic gonadal driver *tj-Gal4* (hereafter tj>*Nup107* KD; Figure. 2B, E). Thus, *Nup107^D364N^* germaria contain elevated numbers of cells with spherical fusomes, characteristic of both GSCs and undifferentiated cystoblasts. Interestingly, an increase in the total number of undifferentiated germ cells is reminiscent of hyperactivation of the Bone Morphogenetic Protein (BMP) signaling pathway (*14*).

**Figure. 2.**
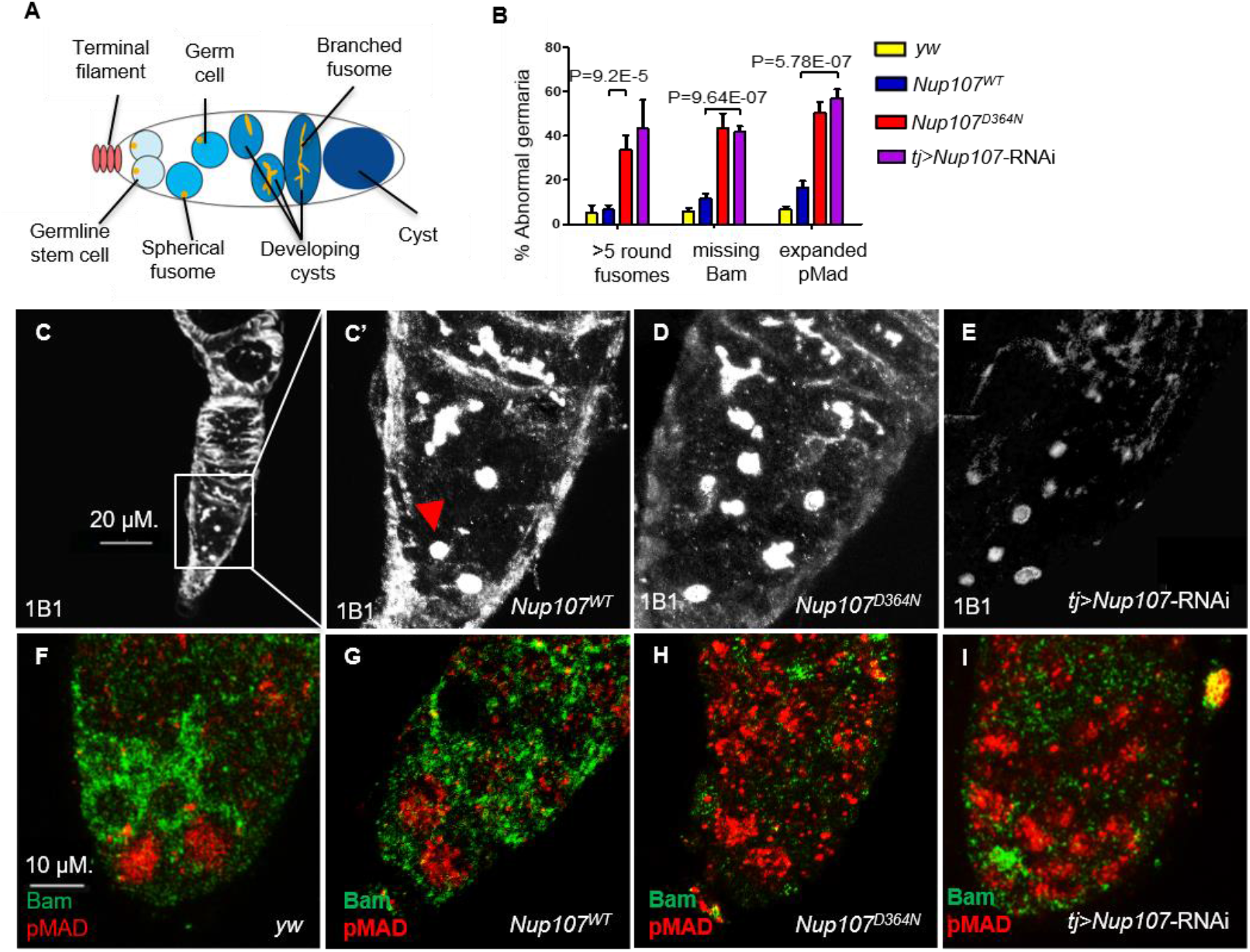
*Nup107^D364N^* adult ovaries demonstrate increased germ line stem cell number. (**A**) A scheme of the *Drosophila* germarium. (**B**) Quantification of spherical fusomes (n=57, 74, 100, 27), Bam (n=127, 137, 139, 64) and pMad (n=122, 121, 225, 48) phenotypes in *yw*, *Nup107^WT^, Nup107^D364N^* and *Nup107* KD, respectively. (**C**) *Nup107^WT^* germaria contain 2-3 cells with spherical fusomes, indicated by arrowhead, while (**D**) *Nup107^D364N^* and (**E**) *tj>Nup107* germaria contain an average of 6 cells with spherical fusomes. (**F**) *yw* and (**G**) *Nup107^WT^* germaria show normal pMad (blue) and Bam (green) expression compared to (**H**) *Nup107^D364N^* and (**I**) *tj>Nup107* germaria which show reduced Bam levels and expanded pMad expression.

We thus wondered if inappropriately expanded BMP signaling is responsible for the failure in GSC differentiation. The somatic terminal filament (TF) and cap cells, which together constitute the GSC niche, normally secrete a BMP ligand, Decapenataplegic (Dpp) (*15, 16*). Binding of Dpp to its cognate receptor Thickveins (Tkv) expressed in GSCs triggers a signal transduction cascade, which ultimately represses the expression of the master differentiation gene *bag of marbles* (*bam*) in the GSC region (*17*). As GSC daughter cells move away from the niche, BMP signaling is weakened, resulting in the induction of Bam expression. We found that Bam expression in mutants was continuously repressed and completely absent in 43% of *Nup107^D364N^* and 42% of *tj>Nup107 KD* stage 2a germaria, compared to 6% and 11% in *yw* and *Nup107^WT^* flies, respectively (Figure. 2B, F-I). As phenotypic consequences resulting from *Nup107* loss seemed analogous to those induced by excess BMP signaling, we sought to analyze the downstream components of the BMP pathway.

The transcriptional response to the BMP signal that emanates from the stem cell niche is mediated by phosphorylation of MAD (pMad) which translocates to the nucleus and with its binding partner, Medea, regulates pathway targets (*17–19*). pMad represses *bam* and thus allows for maintenance of the undifferentiated state and self-renewal of GSCs (*20*). We found that pMad levels in Nup107-compromised germaria are abnormally elevated in regions distant from the GSC niche. Specifically, 50% of *Nup107^D364N^* and 57% of *Nup107* KD germaria show high expression of pMad, compared to 7% of yw and 17% of WT-rescued germaria (Figure. 2B, F-I). Taken together these findings support the notion that compromising Nup107 activity leads to hyperactivation of BMP/Dpp signaling away from the GSC niche and suggest that Nup107 restricts the range and/or strength of Dpp signaling required for proper differentiation of the GSCs.

### Impairment of the GSC progeny differentiation niche in Nup107 mutants

Escort cells (ECs) and their cellular processes, which encapsulate the GSC daughter cells that leave the stem cell niche, constitute a niche that controls germ cell differentiation. This is thought to be achieved, in part, by restricting BMP signaling (*21, 22*). ECs utilize Hh, Wnt, EGFR, and Jak-Stat signaling to prevent BMP signaling in GSC progeny (*23–32*). Many of these signals act to sustain the cytoskeletal structure of the processes that emerge from the ECs. These processes were shown to be essential for the proper differentiation of GSC progeny. The accumulation of undifferentiated GSCs and the expanded range of BMP signaling observed in Nup107 compromised germaria could result from the failure of the EC differentiation niche function. We reasoned that such a failure may be reflected in the cellular extensions emerging from the ECs. To test this idea, ovaries were co-stained with anti-Traffic Jam (TJ) and anti-Coracle (Cora). Cora is a structural protein that is highly expressed in the cellular extensions of ECs (*32, 33*) whereas TJ is a transcription factor expressed in the nuclei of ECs and follicle stem cells in the germarium (*34*). The cellular extensions of the ECs in *yw* and *Nup107^WT^* germaria were readily detected (Figure. 3A, B). In contrast, these cellular extensions were either dramatically reduced or completely lost in over 36% and 48% of the *Nup107^D364N^* and *tj>Nup107 KD* germaria, respectively (Figure. 3C-E). It should be noted that the similar number of escort cells were present in both WT and Nup107 compromised germaria. Together these data imply that compromising Nup107 impairs the formation of cellular extensions in ECs required for restricting the BMP signal in the GSC differentiation niche.

**Figure. 3.**
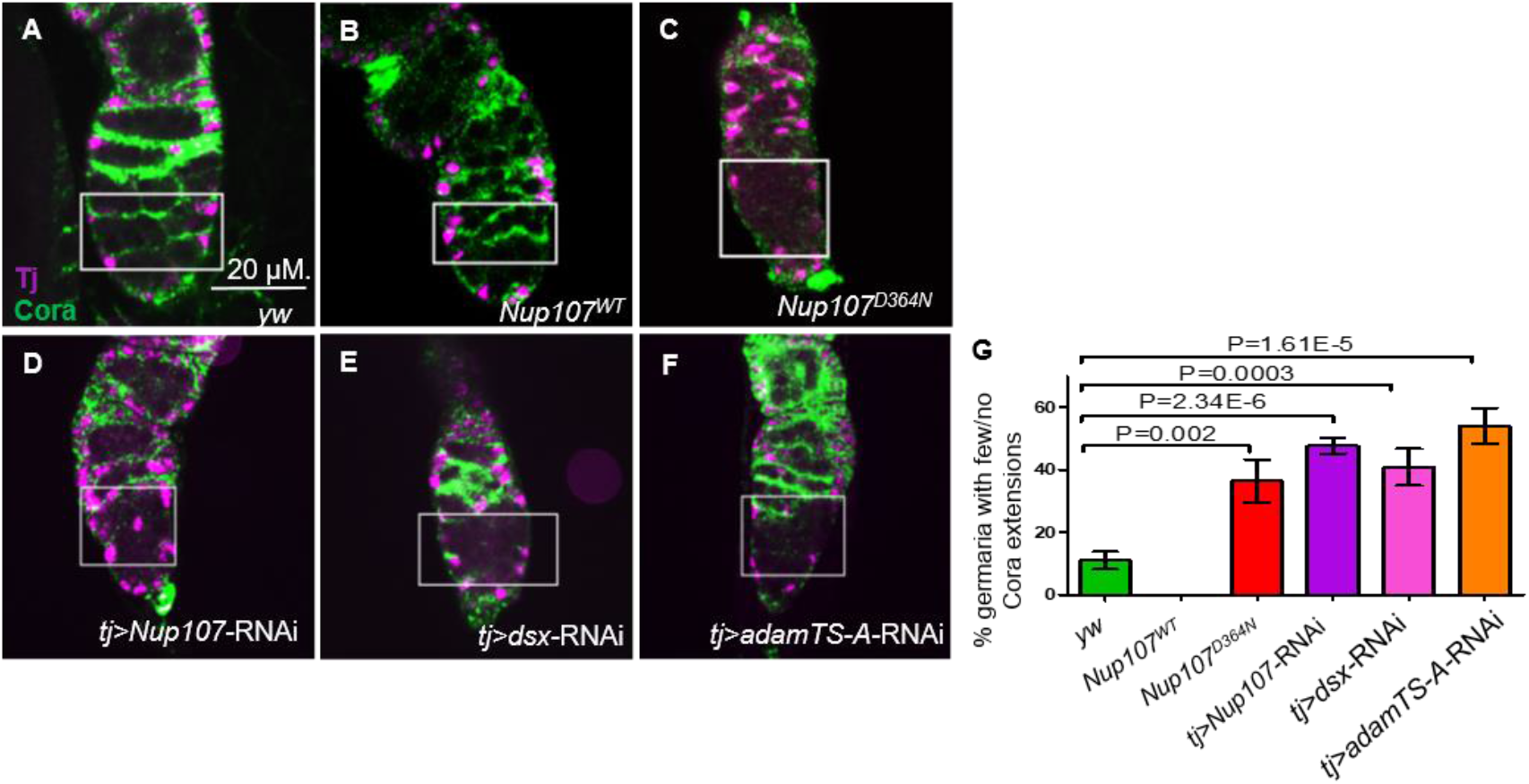
Adult ovaries demonstrate few or no ECs extensions. In germaria taken from (**A**) *yw* and (**B**) *Nup107^WT^*, the ECs extensions were easy to note. In contrast, ECs extensions from (**C**) *Nup107^D364N^*, (**D**) *tj>Nup107 KD,* (**E**) *tj>dsx KD* and (**F**) *tj>adamTS-A KD* germaria were dramatically reduced or lost. (**G**) Quantitation of Coracle extensions from *yw* (n=88), *Nup107^WT^* (n=32), *Nup107^D364N^* (n=25), *tj>Nup107 KD* (n=65)*, tj>dsx KD* (n=60) and *tj>adamTS-A KD (*n=54). Anti-Cora and anti-TJ staining are shown in green and magenta respectively.

### Soma-germline homeostasis in the larval gonad requires Nup107

To trace back the phenotypic consequences of Nup107 loss to the earlier stages of gonad development, we sought to analyze the third instar larval stage (LL3) gonads. At this stage, the *Drosophila* larval gonad consists of a somatically derived stem cell niche and its adjacent primordial germ cells (PGCs, Figure. 4A). Interspersed amongst the PGCs are their somatic support cells, known as Intermingled Cells (ICs). We paid special attention to the ICs as they serve as the progenitors of the ECs. Dissection of *yw*, *Nup107^WT^* and *Nup107^D364N^* larvae showed that their gonads were equally present and readily identifiable in all three genotypes (Figure 3-Figure Supplement 1).

**Figure. 4.**
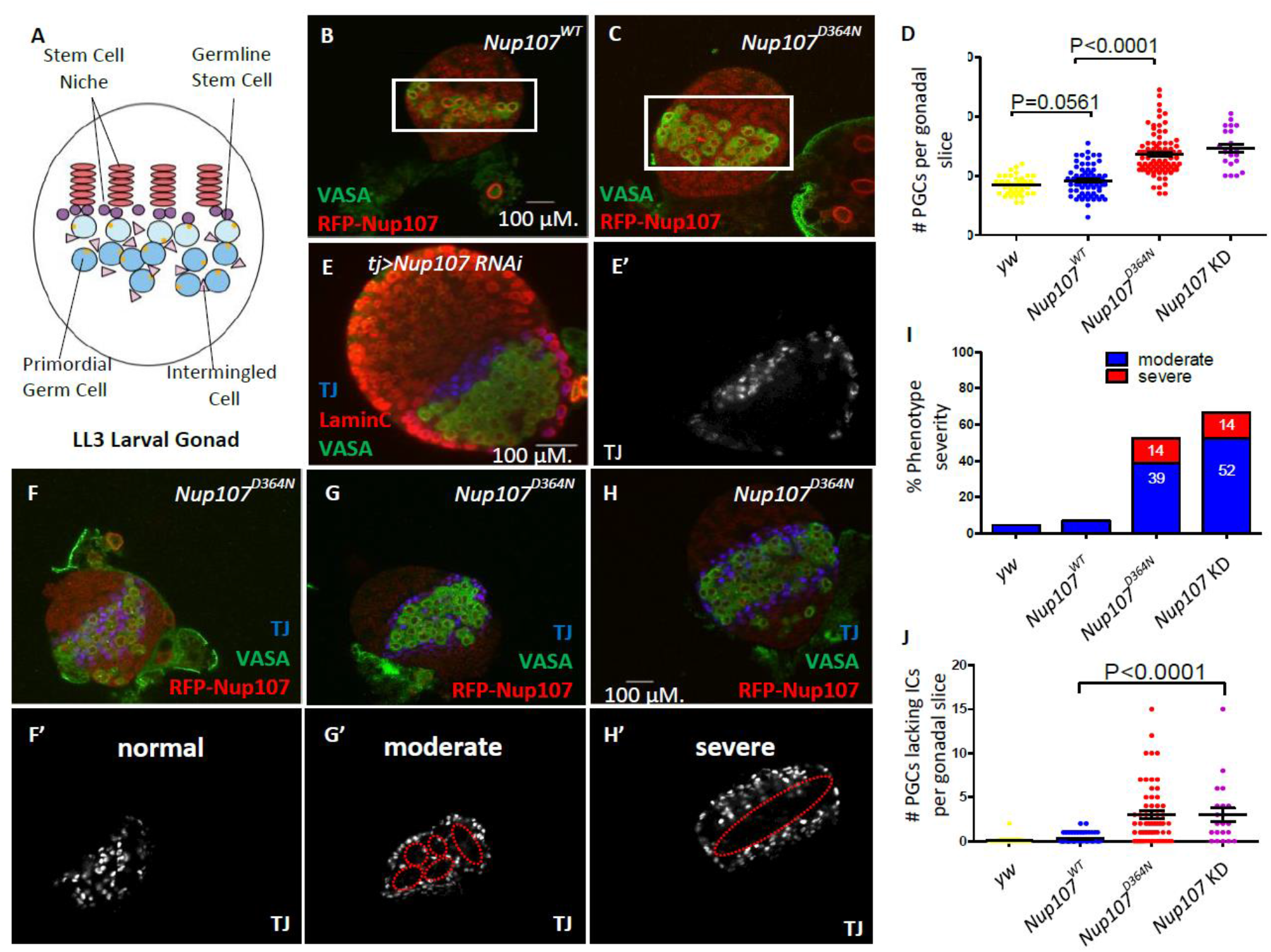
*Nup107^D364N^* larval gonads display aberrant cellular number and arrangement. (**A**) Schematic representation of the LL3 larval gonad with its different cell types. (**B**) Confocal section of *Nup107^WT^* gonad containing on average 18 PGCs (VASA, green), compared to (**C**) 27 in *Nup107^D364N^* gonads. (**D**) Quantitation of the total PGCs per confocal section in each gonad, in *yw*, *Nup107^WT^*, *Nup107^D364N^*, and *Nup107* KD (n=42, 71, 88, 22). (**E**) *Nup107* KD larval gonads contain excess PGCs (VASA, green) and abnormally dispersed ICs (TJ, blue). *Nup107^D364N^* larval gonads exhibit a range of IC dispersion patterns from (**F**) normal to (**G**) moderate to (**H**) severe. (**I**) The percentage of each IC phenotype found in *yw*, *Nup107^WT^*, *Nup107^D364N^* and *Nup107* KD gonads (n=22, 61, 58, 21). (**J**) Representation of the number of PGCs per gonad missing an immediately adjacent IC, due to abnormal dispersion.

To assess if the somatic and germline components of the larval gonad are specified and patterned correctly in a Nup107 compromised background, we stained larval gonads from both *Nup107^D364N^* flies and those with somatic KD of Nup107 (tj>*Nup107* KD), using antibodies which specifically mark the PGCs (VASA) and ICs (TJ). Confocal imaging of these larval gonads revealed that both cell types are present at this developmental stage, however their spatial organization appeared to be disrupted. Moreover, both *Nup107^D364N^* and *tj>Nup107* KD larval gonads contained excess numbers of PGCs (Figure. 4D, E) which formed large clusters devoid of ICs (Figure. 4E’, I, J). While the total number of mutant ICs was unaffected (Figure 4-Figure Supplement 1), these cells failed to intermingle with the PGCs and often remained clustered at the periphery of the gonadal tissue (Figure. 4E-H). Staining with antibodies against an adducin-like molecule (1B1) which marks the fusome, a sub-cellular organelle, revealed elevated numbers of cells with spherical fusomes, normally characteristic of germline stem cells (Figure 3-Figure Supplement 1 B, D). Proper soma-germline communication is necessary for the maintenance of the homeostatic balance required for adequate proliferation and differentiation of both cell types. Altogether these data show that loss of *Nup107* in the somatic gonadal precursors affects the soma-germline communication, adversely affecting the total PGC count and relative positioning of the ICs.

### Sex-determination gene Dsx is a target of Nup107

We thus sought to explore if the Nup107 D364N mutation influences transcription in the female larval gonads and if so, whether the possible changes in the transcriptional profile could be correlated with the developmental defects observed in *Nup107* mutant ovaries. To assess the genome-wide transcriptional changes in the *Nup107* mutant ovaries, we performed an unbiased transcriptome analysis on *yw*, *Nup107^WT^* and *Nup107^D364N^* LL3 larval gonads using RNA-seq (GEO accession number GSE141094). We identified 82 candidate genes (Tables supplement 1, 2) which displayed significant changes in mRNA expression in the larval gonad upon compromising *Nup107*. Among these candidates, we were particularly intrigued by the decreased expression of the DMRT transcription factor family member *doublesex* (Dsx), which is critical for sex-specific differentiation (*35–37*). We confirmed that *dsx* is a target of Nup107 using qRT-PCR (Figure 4-Figure Supplement 2).

*Dsx* is expressed in both sexes, but is regulated via sex-specific alternative splicing, which results in the generation of either female (Dsx^F^) or male (Dsx^M^) specific isoforms (*38–40*). Supporting the conclusion that both Dsx^F^ and Dsx^M^ are determinants of proper sexual development, either loss of individual *dsx* function or simultaneous gain of both *dsx^F^* and *dsx^M^* results in an intersexual phenotype (*41*). Interestingly, despite its crucial role in establishing and maintaining sexually dimorphic differentiation and behavior, *dsx* is expressed in only a select subset of tissues (*42, 43*). Establishing the biological relevance of the localized expression, inactivation of *dsx* using *dsx-Gal4* resulted in reduction in the size of the ovaries (*41*). As *dsx* expression was adversely influenced upon ovary specific loss of Nup107, we sought to test if Dsx plays an important role in female differentiation downstream of Nup107.

To assess this possibility, we first tested if compromising *dsx* in the somatic component of the gonad can mimic the *Nup107^D364N^* phenotypes. Indeed, knockdown of *dsx* expression using somatic driver *tj-Gal4*, resulted in ovarian defects including either partial or complete dysgenesis (in 52% of adult ovaries; Figure. 5A-D). Importantly, compromising *dsx* using *nos-Gal4*, a germline specific driver, did not yield similar phenotypes. Furthermore, *dsx* KD using a *tj-Gal4* driver also led to excess numbers of PGCs as well as a significant number of larval gonads with aberrant IC distribution (Figure. 5E-H). Analysis of germaria compromised for Dsx^F^ function in the ovarian soma largely mimicked the loss of *Nup107* with respect to excess GSCs (distinguishable by the high spherical fusomes; Figure 5I) and loss of the cellular extensions of the ECs (Figure 3E). Importantly, as in the case of *Nup107*, loss of Dsx^F^ in the ovarian soma correlated with aberrant BMP signaling, as reflected in the accumulation of excess nuclear pMad accompanied by the loss of Bam in region 2a of the germaria (Figure. 5J-K).

**Figure. 5.**
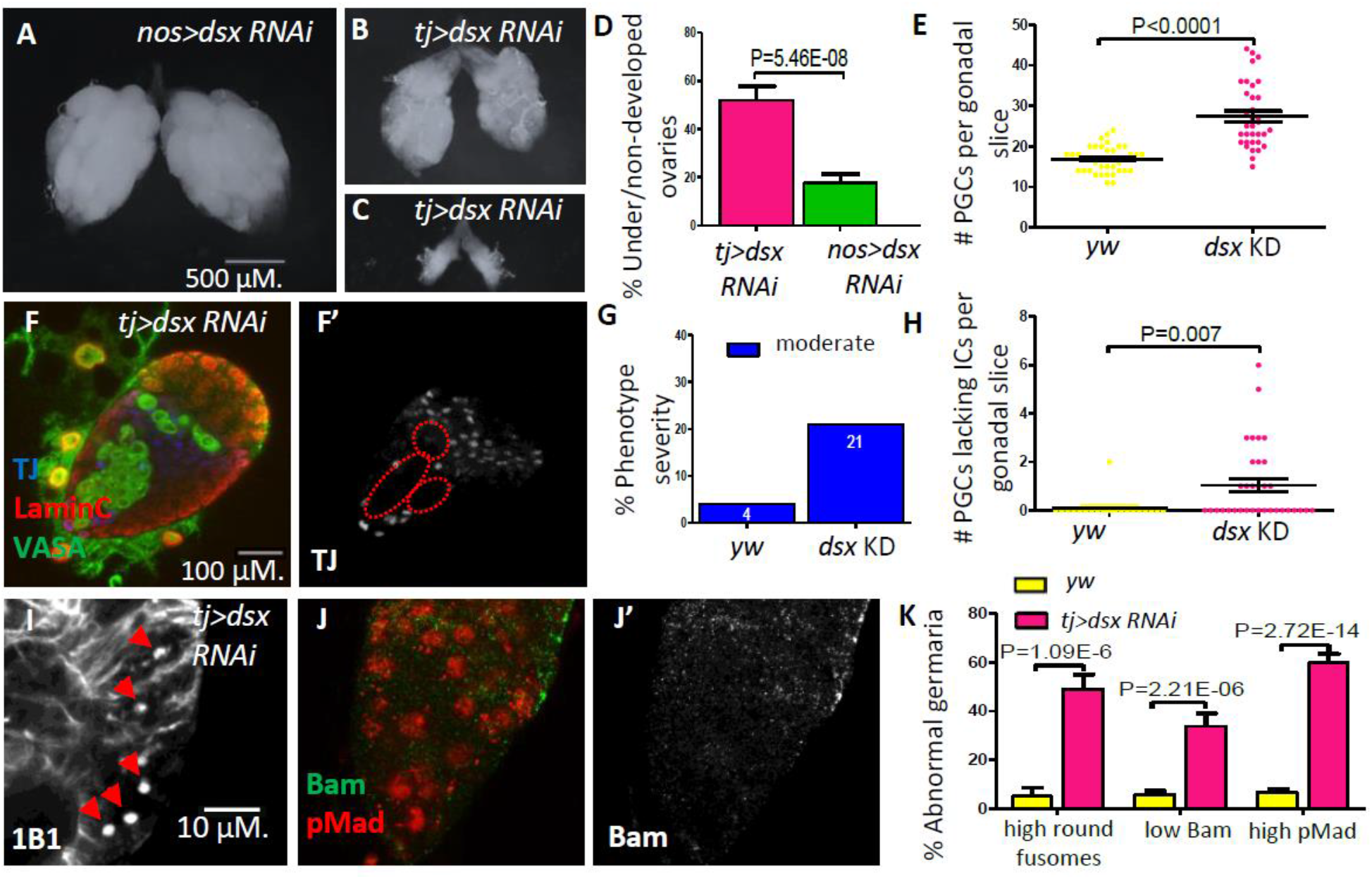
Knockdown of *dsx* in the gonadal soma recapitulates *Nup107^D364N^* phenotypes. (**A**) *dsx* knockdown in the germline (*nos-Gal4*) results in negligible effects, compared to KD in somatic cells (*tj-Gal4*), which results in both (**B**) underdeveloped and (**C**) non-developed ovaries. (**D**) The percentages of under/non-developed ovaries in *tj-Gal4 dsx* KD (n=74) vs *nos-Gal4 dsx* KD (n=100). (**E**) Representation of the number of PGCs per confocal section of each individual gonad in *yw* (n=42) and *dsx* KD (n=34) larvae. (**F**) Larval gonads where *dsx* is knocked down using *tj-Gal4* contain excess PGCs and abnormally dispersed ICs. (**G**) The percentage of IC severity phenotypes found in *yw* (n=22) and *dsx* KD (n=16) larval gonads. (**H**) Quantitation of the number of PGCs per gonad lacking an immediately adjacent IC, as a result of abnormal IC dispersion in *yw* (n=22) and *dsx* KD (n=34) larval gonads. (**I**) *tj*-driven *dsx* KD results in excess spherical fusomes, as well as (**J**) excess pMad (red) expression and missing Bam (green). (**K**) Quantitation of *dsx KD* germaria aberrant phenotypes (n= 53, 119 and 63 respectively).

### Dsx overexpression rescues the phenotypes induced by Nup107 loss

The similarity between the *Nup107* mutant and *dsx* KD ovarian phenotypes prompted us to test whether *dsx* is a critical target of *Nup107*. If so, overexpression of Dsx^F^ alone should mitigate the phenotypes induced by the loss of *Nup107*. We therefore knocked-down *Nup107* and concomitantly overexpressed *Dsx^F^*. Analysis of the resulting ovarian samples revealed that the proportion of underdeveloped ovaries was substantially diminished as compared to both *Nup107^D364N^* and *tj-Gal4* driven *Nup107* KD flies (Figure. 6A-C). The gonads rescued upon simultaneous overexpression of *Dsx^F^* appeared morphologically normal (Figure. 6B), resembling those of control *yw* females. Furthermore, normal patterns and levels of both pMad and Bam expression were restored in region 2a of the germaria (Figure. 6D, E). Taken together these data strongly suggested that Dsx is a key component acting downstream of Nup107 and loss of *Dsx^F^* could account for the stage and sex specific phenotypes associated with compromised Nup107 activity.

**Figure. 6.**
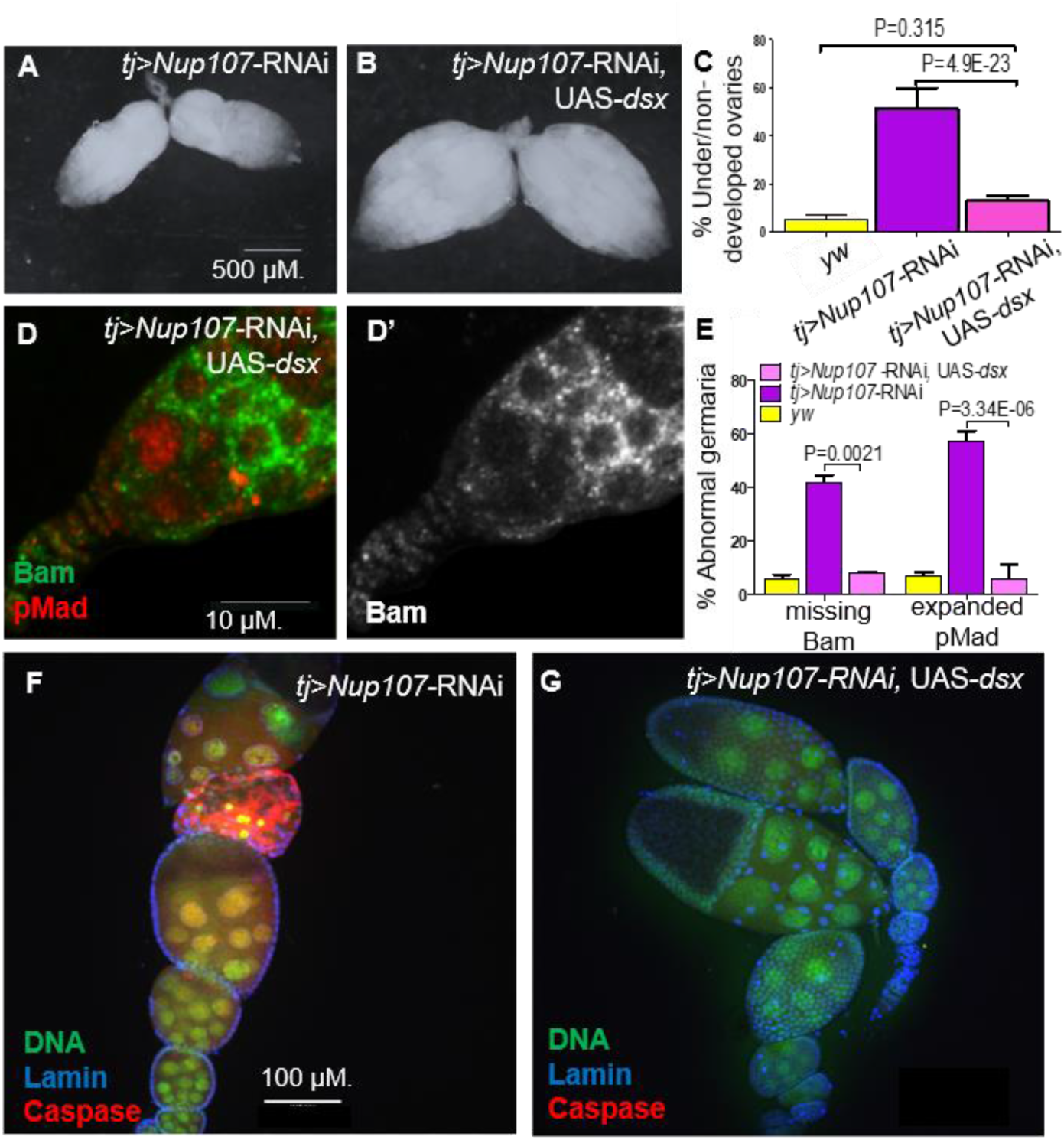
Overexpression of Dsx rescues Nup107 KD ovarian phenotypes. **(A)** RNAi-KD of *Nup107* using *tj-Gal4* driver results in small, underdeveloped ovaries. **(B)** Co-expression of *dsx* with RNAi-KD of Nup107 rescues the underdeveloped phenotype, resulting in normal, robust ovaries. **(C)** Quantitation of under/non-developed ovaries in *yw* (196), *Nup107* KD (n=316) and *Nup107* KD, *dsx* OE flies (n=250). **(D)** *tj-Gal4* driven Nup107 KD, *dsx* OE germarium contains normal pMad (red) expression and **(D’)** normal Bam expression. **(E)** Quantitation of Bam and pMad expression in *yw* (n=127, 122), *Nup107* KD (n=64, 48), and Nup107 KD, *dsx* OE (n=25, 33) germaria. **(F)** 55% of *tj-Gal4* driven *Nup107* KD ovarioles (n=20) showed apoptosis, marked by anti-Caspase3 (red) compared to **(G**) zero *Nup107* KD, *dsx* OE ovarioles (n=15).

The nearly complete rescue observed by the introduction of Dsx^F^ transgene also implied that the two determinants could share important targets. We tested the notion by comparing the 82 genes, identified as Nup107 targets, with previously reported Dsx^F^ targets (*41*). Indeed, the comparison revealed that 47 out of the 82 Nup107 targets (57%, *P*<4.5E-11) were also identified as targets of Dsx (Table supplement 2). In addition to several transcription factors, included in this list are known modulators of BMP signaling as well as multiple components of the extracellular matrix. The substantial overlap between the two lists thus provides the key to the nearly complete rescue. Importantly these data have begun to elucidate how germline-soma communication engineered by the Dsx^F^ can contribute to the establishment of female germline identity.

### BMP modulator AdamTS-A acts downstream of Nup107 and Dsx

Our data demonstrate that that Dsx^F^ regulates the range and/or the strength of BMP signaling from the niche. We were thus intrigued by two other candidate genes identified in our differential transcriptome analysis, which have been previously implicated in modulating the range of BMP signaling. These include AdamTS-A, a metalloprotease that controls extracellular matrix (ECM) assembly. Using qPCR analysis, we found that their expression is appreciably reduced in adult female ovaries compromised for *Nup107* activity as compared to the wild type (Figure. 7G, Figure 7-Figure Supplement 1). Here we have focused our attention on *adamTS-A*, as human ovarian disorders including polycystic ovary syndrome and primary ovarian insufficiency have been correlated with compromised *AdamTS-A* activity (*44–46*). We first examined if the downregulation of *AdamTS-A,* downstream of Nup107, is mediated by Dsx^F^. Supporting the conclusion that transcription of *adamTS-A* is positively regulated by *Dsx^F^*, we observed a significant decrease in the levels of *adamTS-A* transcripts following knockdown of *dsx* (Figure. 7G).

**Figure. 7.**
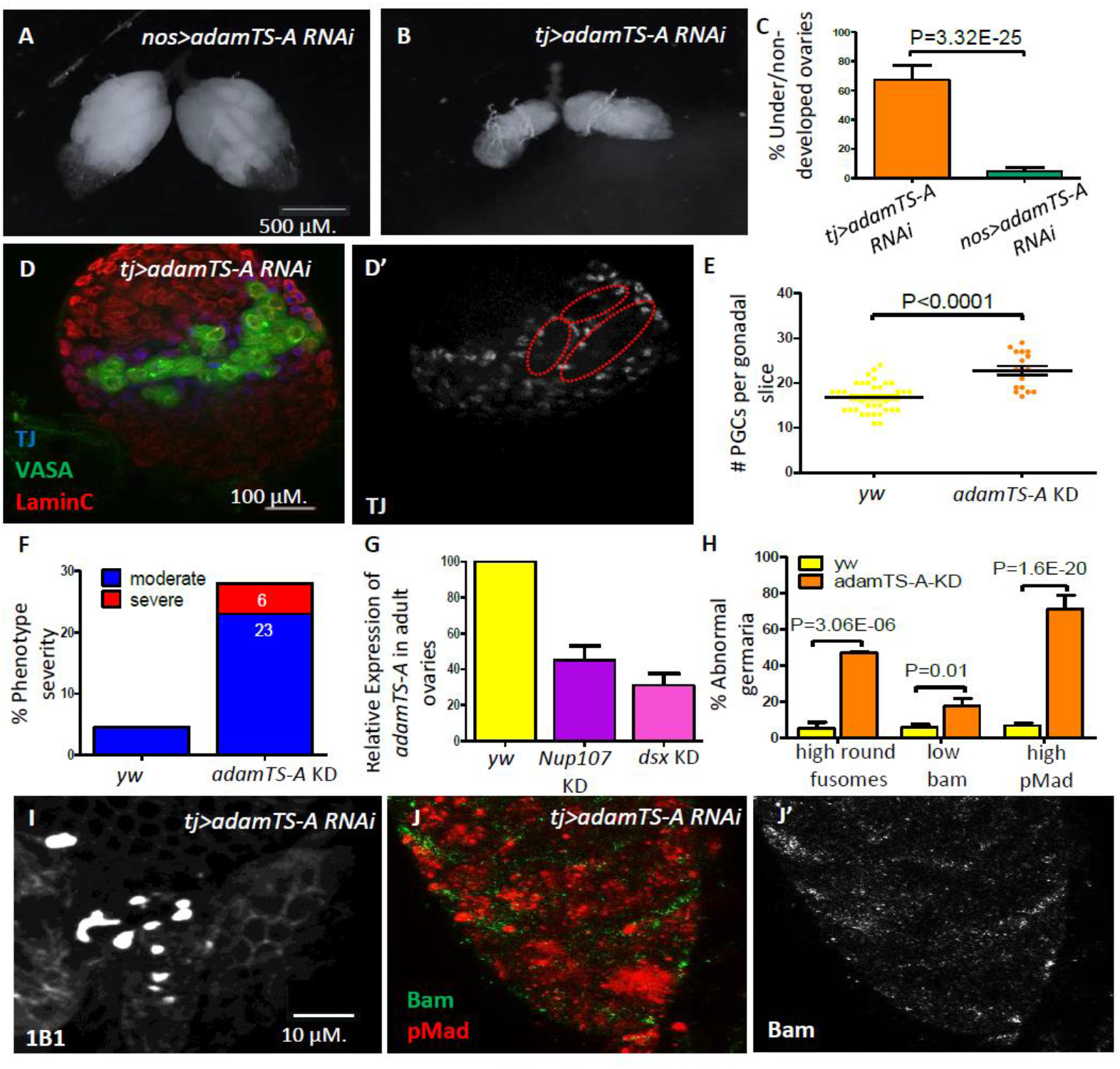
*adamTS-A* KD larval and adult ovaries show aberrant phenotypes. (**A**) Germline KD of *adamTS-A* (*nos-Gal4*) results in negligible effects, compared to somatic KD (*tj-Gal4*), which results in (**B**) severely underdeveloped ovaries. (**C**) Quantitation of under/non-developed ovaries in *tj-* vs *nos-Gal4* driven *adamTS-A* KD (n=126, 108) flies. (**D**) Somatic KD (*tj-Gal4*) of *adamTS-A* results in larval gonads containing excess numbers of PGCs (VASA, green) and abnormally dispersed Intermingled Cells (TJ, blue). (**E**) Quantitation of the total number of PGCs per confocal section in each individual gonad in *yw* (n=42) and *adamTS-A* KD (n=17) larvae. (**F**) The percentage of IC severity phenotypes found in *yw* (n=22) and *adamTS-A* KD (n=17) larval gonads. (**G**) Relative expression of *adamTS-A* measured by RT-qPCR. (**H**) Quantification of cells with round fusomes (n=57), Bam expression (n=145, green), and pMad expression (n=73, red) in *yw and adamTS-A* KD ovaries. *tj-Gal4* driven *AdamTS-A* KD results in (**I**) excess number of cells with spherical fusomes (anti-1B1), (**J**) expanded pMad (red), and (**J’**) reduced Bam expression.

Further, we sought to test if downregulation of *adamTS-A* is an important determinant of ovarian development downstream of Nup107 and/or Dsx^F^. Knockdown of *adamTS-A* in the somatic gonadal cells had severe effects on ovarian development including partial ovarian dysgenesis (Figure. 7B, C), increased number of PGCs (Figure. 7D-F) and GSCs (Figure. 7H, I) in the larval and adult female gonads respectively, with expanded levels of pMad and loss of Bam expression as well as loss of ECs’ extensions in the germarium (Figures. 7H, J and 3F, respectively). In contrast, germline specific knockdown or disruption of its activity using other tissue-specific drivers (wing, eye etc.) resulted in negligible effects (Figure 7A, C, and data not shown). This finding confirmed AdamTS-A as a biologically relevant component that likely acts downstream of both Nup107 and Dsx in this context.

## Discussion

We have shown that Nup107 activity in the somatic component of the gonad is necessary for the proper development of the ICs which enables them to interact with the PGCs. Consistent with the notion, loss of Nup107 affected the behavior of ICs such that these cells showed varying degrees of failure to mingle with the PGCs. A severe failure of ICs and PGCs to interact in the larval gonad is expected to cause ovarian dysgenesis-like phenotype as it is essential for the germarium development and ovariole formation. In the adult ovary, loss of Nup107 impaired the differentiation of the ECs, so that they failed to form the cellular extensions necessary for interactions with the GSCs and regulate their differentiation. The adult ovarian ECs are the descendants of the larval ovarian ICs. Collectively, our results are consistent with the model that Nup107 activity is required in the larval gonad for the specification of the ICs, whereas the adult ovarian phenotypes reflect a secondary consequence of compromising Nup107 activity in the progenitors of the ECs. Our data however do not rule out the possibility that Nup107 function is continuously necessary in different cell populations at different times during ovarian development. Future experiments involving selective knockdown of Nup107 in a temporally restricted manner will shed light on the precise function of Nup107 in this context.

Nevertheless, our studies have revealed that Nup107, a ubiquitously expressed nuclear envelope protein, is a crucial player during female gonad formation. How does an essential housekeeping protein critical for nuclear transport, play such a sex and tissue-specific function? We envisage two possible scenarios, not necessarily mutually exclusive, to explain how the specific mutation in Nup107 results in ovary specific aberrant phenotypes. In the first scenario, Nup107 would specifically mediate nucleocytoplasmic translocation of factor(s) or downstream effector(s) required for ovarian development. Indeed, recent studies have demonstrated that Nup107 is involved in translocation of specific factors. For instance, in the event of DNA-damage, Nup107 directly interacts with the apoptotic protease activating factor 1 (Apaf-1 also known as Drosophila ARK) and mediates its transport into the nucleus to elicit cell-cycle arrest (*47*). Furthermore, it has been shown in tissue culture that specific Nucleoporins, including Nup107, are required for nuclear translocation of SMAD1, an important downstream effector of the Dpp/BMP pathway (*48*).

Alternatively, accumulating evidence has documented that in addition to their primary function in regulating the exchange of molecules between the nucleus and cytoplasm, NPC components may contribute to genome organization and tissue specific regulation of gene expression (*49, 50*) in a nuclear transport-independent manner. Consequently, such moonlighting activities may not be confined to the nuclear envelope which is the primary native location of these proteins. For instance, mammalian Nup107-160 complex (a sub-complex of the NPC of which Nup107 is a key component) has recently been shown to shuttle in and out of GLFG nuclear bodies containing Nup98, a nucleoporin that regulates multiple aspects of gene regulation (*51*). Consistently, Nup107 was shown to regulate levels of specific RNAs through gene imprinting (*52*). Furthermore, using an RNAi-based assay, Nup107 was identified as a positive regulator of OCT4 and NANOG expression in human ESCs (*53*). In this regard, it is noteworthy that the recently published chromatin-binding profile of Nup107 suggested that Nup107 specifically targets active genes (*54*). Altogether these data support the possibility that Nup107 affects transcription of specific target genes in a tissue- and sex- specific manner either directly or indirectly.

In *Drosophila melanogaster*, Sex-lethal (Sxl) is the master determinant of somatic sexual identity, regulating a splicing dependent regulatory cascade resulting in the presence of alternatively spliced sex-specific isoforms of Dsx protein, Dsx-F and Dsx-M, in females and males respectively. Subsequent dimorphic sexual development including sex-specific gonad morphogenesis is under the control of these Dsx isoforms. Consistently, Dsx proteins deploy components of the housekeeping machinery to achieve sex-specific development of the gonads. Thus, such ‘maintenance’ factors are unlikely to be involved in any regulatory capacity. Our data challenge this notion and demonstrate the presence of sexually dimorphic circuitry downstream of a ‘housekeeping’ nuclear envelope protein, Nup107, which regulates the expression the female form of Dsx.

The similar sex-specific and ovary-restricted phenotype associated with compromised Nup107 activity in both humans and flies implies common underlying molecular mechanisms. We have identified Dsx as the primary target acting downstream of Nup107 in *Drosophila* ovarian development. The mammalian homologues of Dsx, the Dmrt family of transcription factors, also function during sex specific gonad development. However, in mammals the main function of *Dmrt* genes in the gonad is to promote male-specific differentiation. While detailed functional analysis is not available, it is plausible that in mammals, another key female-specific transcription factor, like *Foxl2* (female-specific forkhead box L2) may act downstream of Nup107 to substitute for DsxF in flies.

Our observations have also uncovered that Dsx^F^ controls somatic niche function by calibrating the range and/or strength of Dpp/BMP signaling, possibly via modulation of the level and/or activity of the extracellular matrix components. Thus, it will be critical to elucidate how activities of non-sex-specific components such as Nup107 are coordinated with sex-specific regulation to achieve the precise specification and patterning underlying gonad development. This is of particular significance since modulation of BMP signaling circuitry is inextricably linked with the establishment and maintenance of stem cell fate. Importantly, as in the case of Nup107, BMP signaling is also required in a non-sex-specific manner in a variety of developmental contexts. These observations therefore open new avenues towards the critical examination of how a productive molecular dialogue is established between non-sex-specific housekeeping machinery and versatile intersecting developmental pathways, in order to ultimately achieve proper sex-specific gonadogenesis crucial for fertility, and transmission of genetic information.

## Materials and Methods

### Fly Strains

Flies were raised and maintained at 25°C on standard cornmeal yeast extract media. *yw* was used as a wild-type strain. The g*eneration of RFP-Nup107^WT^ and RFP-Nup107^D364N^* transgenic flies was previously described in Weinberg-Shukron et al (*7*). *nanos-Gal4* and *tj-Gal4* were gifted by Lilach Gilboa’s lab (Weizmann Institute of Science, IL). *Nup107* RNAi and *adamTS-A* RNAi lines were provided by the Vienna Drosophila Research Center (VDRC, Vienna, Austria #108047 and #110157). *dsx* RNAi and *dsx* UAS provided by Bloomington Drosophila Stock Center (BDSC; Indiana University; USA; #35645, #41864 and #44223).

The lines’ complete genotypes:

1. *yw: y^1^ w^*^;;*
2. *RFP-Nup107^WT^: y^1^ w^*^; Nup107^E8^/Nup107^E8^*; RFP-Nup107^WT^
3. *RFP-Nup107^D364N^: y^1^ w^*^; Nup107^E8^/Nup107^E8^*; RFP-Nup107^D364N^
4. *nanos-Gal4:* w^*^;; *nos-GAL4::VP16*
5. *tj-Gal4:* w^*^; tj-Gal4/ Cyo;
6. *Nup107* RNAi: y^1^ w^*^; *UAS-Nup107-RNAi*;
7. *adamTS-A* RNAi: w^*^; UAS- *adamTS-A* RNAi;
8. *dsx* RNAi: y^1^; UAS- *dsx –RNAi/Cyo;* or y^1^;; UAS- *dsx -RNAi*
9. UAS *dsx*: y^1^ w^*^;; UAS- *dsx*;

The crosses genotype:

1. *tj-Gal4>Nup107* RNAi:; tj-Gal4/ UAS-*Nup107-RNAi*;
2. *nanos-Gal4>Nup107* RNAi:; *UAS-Nup107-RNAi; nos-GAL4::VP16*
3. *tj-Gal4>adamTS-A* RNAi:; tj-Gal4/ UAS- *adamTS-A* RNAi;
4. *tj-Gal4>dsx* RNAi:; tj-Gal4/ UAS- *dsx –RNAi;* or w; tj-Gal4; UAS- *dsx* - *RNAi*
5. *tj-Gal4>Nup107* RNAi, UAS *dsx*:; tj-Gal4/ UAS-*Nup107-RNAi*; UAS *dsx*

### Adult and Larval Gonad Dissections

Stage LL3 larvae were collected and subsequently dissected for their gonads according to Gilboa and Maimon (*55*). Adult ovaries were dissected from 3-5 day old females placed on yeast for 24-36 hours in the company of males. All experiments were performed at 25°C, and all were independently repeated at least twice. Dissection was performed in Ringer’s solution (130 mM NaCl, 5 mM KCl, 2 mM CaCl2, 50 mM Na2HPO4, pH 7).

### Immunostaining and imaging

Fixation and immunostaining of larval gonads or adult ovaries were carried as previously described by Maimon (*55*) or by Preall (*56*). In brief, gonads or ovaries were fixed in freshly prepared 5% paraformaldehyde (PFA; Electron Microscopy Sciences; Cat# 15714) for 30 min at room temperature. Blocking was carried out in wash buffer supplemented with 1% bovine serum albumin (BSA; MP Biomedicals; Cat. #160069). Primary antibodies were diluted and incubated overnight at 4°C in wash buffer supplemented with 0.3 % BSA. The following primary antibodies were used: guinea-pig anti-Tj (1:10,000; gifted by Dorothea Godt’s lab at the University of Toronto, Toronto, CA), rat IgM anti-VASA (1:100; Developmental Studies Hybridoma Bank (DSHB; Iowa City, IA, USA)), mouse anti-Hts (1B1; 1:20; DSHB), mouse anti-Bam (1:50; DSHB), mouse anti-Cora (C566.9; 1:100; DSHB), rabbit anti-smad3 (1:100; abcam, Cambridge, MA, USA; #ab52903), rabbit anti-cleaved caspase3 (1:200; Cell Signaling Technology, CST, Danvers, MA, USA; #9661) and mouse anti Lamin C (1:20; gifted by Yosef Gruenbaum’s lab at the Hebrew University of Jerusalem, IL). Secondary antibodies (1:400) were conjugated to either Cy2, Cy3 or Cy5 (Jackson Immuno Research Laboratories; West Grove, PA, USA). Ovaries were mounted in Vectashield (Vector Laboratories; Burlingame, CA, USAVE-H-1000). Images were taken on a TE2000-E confocal microscope (Nikon) using x20 or x60 objectives, occasionally with an additional x1.5 zoom. Figures were edited using Adobe Photoshop CC 2017.

### Quantitative real-time PCR analysis

Total RNA was isolated from larval gonads at stage LL3 (~80 per sample) of *yw, Nup107^WT^* and *Nup107^D364N^* using the RNeasy mini kit (Qiagen Valencia, CA, USA; #74104). Briefly, wandering third instar larvae were collected, females were selected and dissected into tubes in liquid nitrogen and lysis buffer was added. Total RNA was extracted as per the manufacturer’s instructions. cDNA was made using the high-capacity cDNA reverse transcription kit (Applied Biosystems, Foster City, CA, USA) using an equal amount of total RNA from each sample. Real-time q-PCR analyses were carried out using the Powersyber Green PCR Master Mix and QuantStudio 12k flex (Applied Biosystems). Rsp17 and TBP2 served as reference genes using the comparative Ct method. Each sample was analyzed in triplicate; results were confirmed by at least two independent experiments. Primer sequences (from HyLabs, Israel, LTD) used for qPCR were:

dm_Doublesex_F, 5′-TTGCCGATCTCAGTTTCCGT-3′;

dm_Doublesex_R, 5′- GCTCCCAAGGATAGCGGAAT-3′;

dm_AdamTS-A_F, 5′- GGGAATGAGCCGAACAAGAC-3′;

dm_AdamTS-A_R, 5′- AAGTTCTGGTCGGGATAGCC-3′.

### Statistical Analysis

The number of under/non-developed adult ovaries in wild-type and mutant flies, the varied expression of 1B1, bam, Cora, as well as the number of spherical fusomes in germaria were compared pairwise using Fisher’s Exact Test for 2×2 tables. The raw 2-tail p-values were adjusted for the multiple comparisons using either the Bonferroni correction or Holm’s modification (*57*) thereof, as appropriate. P values of less than 0.05 were considered significant.

The number of Primordial Germ Cells, Intermingled Cells and fusomes in groups of larval gonads were compared using a Kruskal-Wallis Test (*58*). In experiments where the differences among the groups were found to be significant (K-W p-value <0.05), pairwise comparisons were carried out using Conover’s post-hoc test (*59*).

The statistical significance of the observed overlap between our Nup107 target genes list and the previously reported Dsx targets was calculated using the hypergeometric test (https://systems.crump.ucla.edu/hypergeometric/index.php). The specific parameters were as follows: number of successes k=47; sample size s=82; number of successes in the population M=3717; population size N=15835. The results obtained were: expected number of successes = 19.2481212503947. The results are enriched 2.44 fold compared to expectations hypergeometric p-value = 4.4742336544062e-11.

### RNA-Seq

Total RNA was isolated from *yw*, *RFP-Nup107^WT^*, and *RFP-Nup107^D364N^ Drosophila* larval gonads following dissection at stage LL3 using the RNeasy Kit (Qiagen) according to the manufacturer’s protocol. RNA purity and concentration were determined by T-042 NanoDrop Spectrophotometer (Thermo Fisher Scientific Inc., Waltham, MA, USA) and integrity by 2100 Bio-analyzer (Agilent Technologies, CA, USA). Total RNA was reverse transcribed to cDNA using SENSE Total RNA-Seq Library Prep Kit for Illumina (Lexogen, Vienna, Austria), according to the manufacturer’s protocol, with poly-A selection. Libraries were multiplexed and sequenced on Illumina’s NextSeq 500 machine, with a configuration of 75 cycles, single read. Raw reads were processed to remove low quality, error prone and adapter sequences, according to Lexogen’s SENSE libraries recommendations. High quality reads were aligned to the fly genome, assembly BDGP6, that was supplemented with the sequences of GFP and the RFP-Nup constructs. Alignment was performed with TopHat, allowing for up to 5 mismatches per read. Differential expression analysis, for all genes from release 84 of the Ensembl database, was performed with the DESeq2 package, using default parameters, including the threshold for significance that was padj < 0.1. Significant genes were further filtered to include only genes whose up- or down-regulation was greater in the mut/yw comparison than in the wt/yw one. For that end, a difference of at least 0.1 between the absolute log2FoldChange(mut/yw) and absolute log2FoldChange(wt/yw) was used as the filtering threshold. All raw data, as well as software versions and parameters, have been deposited in NCBI’s Gene Expression Omnibus (*60*) and are accessible through GEO Series accession number GSE141094.

## Acknowledgments

We thank L. Gilboa, the Developmental Studies Hybridoma Bank, the Vienna Drosophila RNAi Center, and the Bloomington Drosophila Stock Center for reagents and fly stocks. We thank Z. Paroush for critical reading and comments on the manuscript. We thank N. Grover for assistance with statistical analysis. We also thank Mira Korner, Michal Bronstein, and Temima Schnitzer-Perlman at the Center for Genomic Technologies at the Hebrew University of Jerusalem for their assistance and expertise in RNA-seq.

## Abbreviations used in this paper

AdamTS-A: A disintegrin-like and metalloprotease domain [reprolysin type] with thrombospondin type 1 motifs
*bam*: *bag of marbles*
BMP: Bone Morphogenetic Protein
Cora: Coracle
Dpp: Decapenataplegic
Dsx: Doublesex
ECM: extracellular matrix
ECs: Escort cells
*Foxl2*: female-specific forkhead box L2
GSC: germline stem cells
ICs: Intermingled Cells
KD: knockdown (KD)
LL3: late third instar larval stage
*Nup107*: nucleoporin-107
*nos-Gal4*: *nanos-Gal4*
PGCs: primordial germ cells
pMad: phosphorylated Mothers against dpp
TF: terminal filament
Sxl: Sex-lethal
Tkv: Thickveins
*tj-Gal4*: *traffic jam-Gal4*
XX-OD: XX ovarian dysgenesis

## Author contributions

T.S. and T.L. designed, performed, and analyzed the majority of the experiments, with R.K., A.D, and D.R. being involved in the investigation and analysis at earlier stages; and M.Y.G. and S.L. being involved in the later stages. T.S. performed all gonadal analyses while T.L. performed germarium experiments. A.W.S. performed the early stages of the molecular work. Experiments in the wing and eye discs were carried out by T.B. and V.M. T.S. and D.R. extracted the gonadal tissue for the RNA-Seq. RNA-Seq bioinformatic analysis was performed by Y.N. Funding was acquired by O.G., D.Z., and G.D. The project was conceptualized by O.G and G.D. The original manuscript was written by T.S., O.G. and G.D., and subsequently reviewed and edited by T.L and M.Y.G., O.G. supervised the project.

## Data and materials availability

All raw RNA-seq data, as well as software versions and parameters, have been deposited in NCBI’s Gene Expression Omnibus and are accessible through GEO Series accession number GSE141094. All other data is available in the main text or the supplementary materials.

## Supplementary Files

### Supplementary Figure legends

**Figure 3-Figure supplement 1. Female larval gonads compromised for *Nup107* show characteristic aberrations.** (**A**) Unlike their adult counterparts, the *Nup107^D364N^* female larval gonads are present at LL3. (**B**) The difference between the total number of cells with spherical fusomes in *yw* (n=11) *Nup107^WT^* (n=13) and *Nup107^D364N^* (n=17) larval gonads is significant. (**C**) Similar numbers of cells with round fusomes (anti-1B1) can be seen *in yw* and *Nup107^WT^* gonads, while being substantially elevated in (**D**) *Nup107^D364N^* gonads.

**Figure 4-Figure supplement 1. Total number of Intermingled Cells (ICs) remains unaffected in the larval gonads compromised for *Nup107*.** (**A**) The difference between the average number of ICs in *yw* (n=13), *Nup107^WT^* (n=14) and *Nup107^D364N^* (n=12) larval gonads is statistically insignificant. P value represents comparison of all three groups.

**Figure 4-Figure supplement 2. Reduction in *dsx* and *adamTS-A* transcription upon compromising Nup107 activity in the *Drosophila* larval gonad.** (**A**) Quantitative real-time PCR based validation was performed by normalizing the relative expression levels to the abundance of housekeeping gene RpS17.

**Figure 7-Figure supplement 1. *Collagen type IV alpha 1* (*col4a1*) expression levels are reduced in both *Nup107* KD and *dsx* KD adult ovaries.** (**A**) Relative expression of *col4a1* in the *Drosophila* adult ovary, measured by RT-qPCR. RpS17 and TBP2 expression were used as controls. Statistical significance was calculated using the comparative Ct method.

### Supplementary tables

**Table supplement 1.** Processed data results of transcriptomic analysis performed on *Nup107^WT^* and *Nup107^D364N^* larval gonads.

**Table supplement 2.** List of 82 candidate genes with differential expression following Nup107 loss identified in transcriptomic analysis, with emphasis on those found to be *dsx* targets.

## References

1. M. Van Doren. (Springer, New York, NY, Molecular Biology Intelligence Unit, 2006).

2. S. Staab, A. Heller, M. Steinmann-Zwicky, Somatic sex-determining signals act on XX germ cells in Drosophila embryos. Development 122, 4065 (1996).

3. M. Wawersik et al., Somatic control of germline sexual development is mediated by the JAK/STAT pathway. Nature 436, 563–567 (2005).

4. T. DeFalco, N. Camara, S. Le Bras, M. Van Doren, Nonautonomous sex determination controls sexually dimorphic development of the Drosophila gonad. Dev Cell 14, 275–286 (2008).

5. R. Nothiger, M. Jonglez, M. Leuthold, P. Meier-Gerschwiler, T. Weber, Sex determination in the germ line of Drosophila depends on genetic signals and inductive somatic factors. Development 107, 505 (1989).

6. I. A. Hughes, Disorders of sex development: a new definition and classification. Best Pract Res Clin Endocrinol Metab 22, 119–134 (2008).

7. A. Weinberg-Shukron et al., A mutation in the nucleoporin-107 gene causes XX gonadal dysgenesis. J Clin Invest 125, 4295–4304 (2015).

8. Y. Ren et al., Functional study of a novel missense single-nucleotide variant of Nup107 in two daughters of Mexican origin with premature ovarian insufficiency. Mol Genet Genomic Med 6, 276–281 (2018).

9. T. Levi, A. Sloutskin, R. Kalifa, T. Juven-Gershon, O. Gerlitz, Efficient In Vivo Introduction of Point Mutations Using ssODN and a Co-CRISPR Approach. Biol Proced Online 22, 14 (2020).

10. R. C. King. (Academic Press, New York, 1970).

11. A. Spradling. (Cold Spring Harbor Lab Press, New York, 1993).

12. M. de Cuevas, A. C. Spradling, Morphogenesis of the Drosophila fusome and its implications for oocyte specification. Development 125, 2781–2789 (1998).

13. H. Lin, A. C. Spradling, Fusome asymmetry and oocyte determination in Drosophila. Dev Genet 16, 6–12 (1995).

14. C.-H. Zhu, T. Xie, Clonal expansion of ovarian germline stem cells during niche formation in Drosophila. Development 130, 2579 (2003).

15. T. Xie, A. C. Spradling, decapentaplegic is essential for the maintenance and division of germline stem cells in the Drosophila ovary. Cell 94, 251–260 (1998).

16. T. Xie, A. C. Spradling, A niche maintaining germ line stem cells in the Drosophila ovary. Science 290, 328–330 (2000).

17. D. McKearin, B. Ohlstein, A role for the Drosophila bag-of-marbles protein in the differentiation of cystoblasts from germline stem cells. Development 121, 2937–2947 (1995).

18. X. H. Feng, R. Derynck, Specificity and versatility in tgf-beta signaling through Smads. Annu Rev Cell Dev Biol 21, 659–693 (2005).

19. B. Schmierer, C. S. Hill, TGFbeta-SMAD signal transduction: molecular specificity and functional flexibility. Nat Rev Mol Cell Biol 8, 970–982 (2007).

20. X. Song et al., Bmp signals from niche cells directly repress transcription of a differentiation-promoting gene, bag of marbles, in germline stem cells in the Drosophila ovary. Development 131, 1353–1364 (2004).

21. D. Kirilly, S. Wang, T. Xie, Self-maintained escort cells form a germline stem cell differentiation niche. Development 138, 5087–5097 (2011).

22. X. Wang et al., Histone H3K9 trimethylase Eggless controls germline stem cell maintenance and differentiation. PLoS Genet 7, e1002426 (2011).

23. X. Wang, A. Page-McCaw, Wnt6 maintains anterior escort cells as an integral component of the germline stem cell niche. Development 145, (2018).

24. C. Y. Tseng et al., Smad-Independent BMP Signaling in Somatic Cells Limits the Size of the Germline Stem Cell Pool. Stem Cell Reports 11, 811–827 (2018).

25. T. Lu et al., COP9-Hedgehog axis regulates the function of the germline stem cell progeny differentiation niche in the Drosophila ovary. Development 142, 4242–4252 (2015).

26. J. Huang, A. Reilein, D. Kalderon, Yorkie and Hedgehog independently restrict BMP production in escort cells to permit germline differentiation in the. Development 144, 2584–2594 (2017).

27. V. I. Mottier-Pavie, V. Palacios, S. Eliazer, S. Scoggin, M. Buszczak, The Wnt pathway limits BMP signaling outside of the germline stem cell niche in Drosophila ovaries. Dev Biol 417, 50–62 (2016).

28. S. Wang et al., Wnt signaling-mediated redox regulation maintains the germ line stem cell differentiation niche. Elife 4, e08174 (2015).

29. L. Luo, H. Wang, C. Fan, S. Liu, Y. Cai, Wnt ligands regulate Tkv expression to constrain Dpp activity in the Drosophila ovarian stem cell niche. J Cell Biol 209, 595–608 (2015).

30. N. Hamada-Kawaguchi, B. F. Nore, Y. Kuwada, C. I. Smith, D. Yamamoto, Btk29A promotes Wnt4 signaling in the niche to terminate germ cell proliferation in Drosophila. Science 343, 294–297 (2014).

31. M. Upadhyay et al., Transposon Dysregulation Modulates dWnt4 Signaling to Control Germline Stem Cell Differentiation in Drosophila. PLoS Genet 12, e1005918 (2016).

32. I. Maimon, M. Popliker, L. Gilboa, Without children is required for Stat-mediated zfh1 transcription and for germline stem cell differentiation. Development 141, 2602–2610 (2014).

33. M. J. Fairchild, C. M. Smendziuk, G. Tanentzapf, A somatic permeability barrier around the germline is essential for Drosophila spermatogenesis. Development 142, 268–281 (2015).

34. M. A. Li, J. D. Alls, R. M. Avancini, K. Koo, D. Godt, The large Maf factor Traffic Jam controls gonad morphogenesis in Drosophila. Nat Cell Biol 5, 994–1000 (2003).

35. C. K. Matson, D. Zarkower, Sex and the singular DM domain: insights into sexual regulation, evolution and plasticity. Nat Rev Genet 13, 163–174 (2012).

36. P. E. Hildreth, doublesex, recessive gene that transforms both males and females of Drosophila into intersexes. Genetics 51, 659–678 (1965).

37. B. S. Baker, K. A. Ridge, Sex and the single cell. I. On the action of major loci affecting sex determination in Drosophila melanogaster. Genetics 94, 383–423 (1980).

38. K. C. Burtis, K. T. Coschigano, B. S. Baker, P. C. Wensink, The doublesex proteins of Drosophila melanogaster bind directly to a sex-specific yolk protein gene enhancer. Embo j 10, 2577–2582 (1991).

39. B. S. Baker, M. F. Wolfner, A molecular analysis of doublesex, a bifunctional gene that controls both male and female sexual differentiation in Drosophila melanogaster. Genes Dev 2, 477–489 (1988).

40. K. C. Burtis, B. S. Baker, Drosophila doublesex gene controls somatic sexual differentiation by producing alternatively spliced mRNAs encoding related sex-specific polypeptides. Cell 56, 997–1010 (1989).

41. E. Clough et al., Sex- and tissue-specific functions of Drosophila doublesex transcription factor target genes. Dev Cell 31, 761–773 (2014).

42. L. E. Sanders, M. N. Arbeitman, Doublesex establishes sexual dimorphism in the Drosophila central nervous system in an isoform-dependent manner by directing cell number. Dev Biol 320, 378–390 (2008).

43. Y. Yang, W. Zhang, J. R. Bayrer, M. A. Weiss, Doublesex and the regulation of sexual dimorphism in Drosophila melanogaster: structure, function, and mutagenesis of a female-specific domain. J Biol Chem 283, 7280–7292 (2008).

44. S. Özler et al., Role of Versican and ADAMTS-1 in Polycystic Ovary Syndrome. Journal of clinical research in pediatric endocrinology 9, 24–30 (2017).

45. D. L. Russell, H. M. Brown, K. R. Dunning, ADAMTS proteases in fertility. Matrix Biol 44-46, 54–63 (2015).

46. E. A. Knauff et al., Genome-wide association study in premature ovarian failure patients suggests ADAMTS19 as a possible candidate gene. Hum Reprod 24, 2372–2378 (2009).

47. L. Jagot-Lacoussiere et al., DNA damage-induced nuclear translocation of Apaf-1 is mediated by nucleoporin Nup107. Cell Cycle 14, 1242–1251 (2015).

48. X. Chen, L. Xu, Specific nucleoporin requirement for Smad nuclear translocation. Mol Cell Biol 30, 4022–4034 (2010).

49. M. Raices, M. A. D’Angelo, Nuclear pore complexes and regulation of gene expression. Curr Opin Cell Biol 46, 26–32 (2017).

50. M. A. D’Angelo, Nuclear pore complexes as hubs for gene regulation. Nucleus 9, 142–148 (2018).

51. S. Morchoisne-Bolhy et al., Intranuclear dynamics of the Nup107-160 complex. Mol Biol Cell 26, 2343–2356 (2015).

52. S. S. Sachani et al., Nucleoporin 107, 62 and 153 mediate Kcnq1ot1 imprinted domain regulation in extraembryonic endoderm stem cells. Nature Communications 9, 2795 (2018).

53. P. Onal et al., Gene expression of pluripotency determinants is conserved between mammalian and planarian stem cells. Embo j 31, 2755–2769 (2012).

54. A. Gozalo et al., Core Components of the Nuclear Pore Bind Distinct States of Chromatin and Contribute to Polycomb Repression. Molecular Cell 77, 67–81.e67 (2020).

55. Maimon, I. & Gilboa, L. Dissection and staining of Drosophila larval ovaries. J Vis Exp, doi:10.3791/2537 (2011).

56. Preall, J. B., Czech, B., Guzzardo, P. M., Muerdter, F. & Hannon, G. J. shutdown is a component of the Drosophila piRNA biogenesis machinery. Rna 18, 1446–1457, doi:10.1261/rna.034405.112 (2012).

57. Edgar, R., Domrachev, M. & Lash, A. E. Gene Expression Omnibus: NCBI gene expression and hybridization array data repository. Nucleic Acids Res 30, 207–210, doi:10.1093/nar/30.1.207 (2002).

58. Holm, S. A simple sequentially rejective multiple test procedure. Scand. J. Statist. 6, 65–70 (1979).

59. Conover, W. J. Practical nonparametric statistics. Wiley, NY, 3rd Ed, §5.2 (1999).

60 Kruskal-Wallis. Inference for categorical data, singly ordered RcX table, StatXact. Cytel Inc, Cambridge MA., version 10 §21.5 (2013).

